# “A Novel Therapeutic Approach to Corneal Alkaline Burn Model by Targeting Fidgetin-like 2, a Microtubule Regulator”

**DOI:** 10.1101/2020.06.18.159533

**Authors:** Jessie Wang, Abhinav Dey, Adam Kramer, Yuan Miao, Juan Liu, Lisa Baker, Joel Friedman, Parimala Nacharaju, Roy Chuck, Cheng Zhang, David J. Sharp

## Abstract

**Purpose:** To determine the efficacy of nanoparticle-encapsulated FL2 siRNA (FL2-NPsi), a novel therapeutic agent targeting the Fidgetin-like 2 (FL2) gene, for the treatment of corneal alkaline chemical injury.

**Methods:** Eighty 12-week-old, male Sprague-Dawley rats were divided evenly into 8 treatment groups: prednisolone, empty nanoparticles, control-NPsi (1 μM, 10 μM, 20 μM) and FL2-NPsi (1 μM, 10 μM, 20 μM). An alkaline burn was induced onto the cornea of each rat, which was then treated for 14 days according to group assignment. Clinical (N=10 per group), histopathologic (N=6 per group), and immunohistochemical (N=4 per group) analyses were conducted to assess for wound healing. FL2-NPsi-mediated knockdown of FL2 was confirmed by *in vitro* qPCR. Toxicity assays were performed to assess for apoptosis (TUNEL assay, N=3 per group) and nerve damage (whole mount immunochemical staining, N=2 per group). Statistical analyses were performed using student’s t-test and ANOVA.

**Results:** Compared to controls, FL2-NPsi-treated groups demonstrated enhanced corneal wound healing, with the 10 and 20 μM FL2-NPsi-treated groups demonstrating maximum rates of corneal re-epithelialization (p=0.0003 at Day 4 and p<0.0001 at Day 8) as assessed by ImageJ software, enhanced corneal transparency, and improved stromal organization on histology. Immunohistochemical analysis of vascular endothelial cells, macrophages, and neutrophils did not show significant differences between treatment groups. FL2-NPsi was not found to be toxic to nerves or induce apoptosis (p=0.917).

**Conclusion:** Dose-response studies found both 10 and 20 μM FL2-NPsi to be efficacious in this rat model. FL2-NPsi may offer a novel treatment for corneal alkaline chemical injuries.

## Introduction

There are an estimated 1.8 million cases of ocular trauma in the US every year, with burn wounds accounting for 7-18% of eye injuries presenting to emergency rooms, 84% of which are due to chemical injuries [1]. Alkaline chemical injuries oftentimes lead to poor visual outcomes due to delayed corneal re-epithelialization, persistent inflammation, and corneal ulceration and scarring [2]. Typically, they are treated with extensive flushing of the eye and topical application of corticosteroids, which target the early stages of inflammation, but have no effect on epithelial migration and interfere with corneal stroma repair, making the tradeoff often undesirable [3].

The corneal epithelium is the outermost layer of cells of the eye and plays an essential role in maintaining the smoothness of the optical surface and the health of the corneal stroma [4]. The corneal epithelium is largely maintained through the production of cells by stem cells in the limbal corneal region. During homeostasis, centripetal migration of these newly formed corneal epithelial cells moves cells through multiple corneal zones into the central cornea, turning over the epithelium every 7 to 10 days [5]. In instances of small injuries to the cornea, adjacent epithelial cells simply spread to fill the defects [6]. Models of corneal scrapes and abrasions produce epithelial defects that are completely healed within 24-72 hours, a time course often too short for detailed comparative analysis of wound healing. However, larger corneal wounds (e.g. burns, severe trauma) cannot be covered by adjacent cells and instead require increased production and extended migration of large numbers of corneal epithelial cells from the limbus into the wound zone. Accordingly, these wounds take longer to heal, offering a valuable model for ocular wound healing studies [7–9].

Current therapeutics for corneal inflammatory and epithelial wound healing have focused on amniotic membranes, topical steroids, and extracellular signaling factors, with limited success [10]. Small interfering RNAs (siRNA) are non-coding RNA molecules between 20-25 nucleotide in length, which interfere with the expression of target genes with complementary nucleotide sequences by degrading mRNA after transcription, preventing translation. This occurs within the RNA interference (RNAi) pathway within the cell. While small-interfering RNA (siRNA)-based therapies have shown some promise for treating a wide range of maladies, including wound closure [11–13], the effective delivery of therapeutic siRNA has proven difficult in many circumstances. siRNA is a relatively large and negatively-charged molecule, making it impermeable to cellular membranes. However, the development and refinement of siRNA delivery systems with nanoparticles is helping to circumvent this challenge [14, 15]. While previous siRNA studies have shown limited success [16, 17], the combination of new delivery platforms and the appropriate siRNA gene target could result in new ocular therapies.

Multiple investigations in recent years have looked into harnessing the intracellular cytoskeleton, particularly microtubules, for corneal wound healing. Hollow polymeric filaments composed of tubulin subunits, microtubules provide structural support for the cell and are an important substrate for many of the molecular motor proteins responsible for intracellular transport. Regulation of the dynamic properties of microtubules are critically important for the capacity of cells to close wounds, especially near the cell periphery. Experimental alteration of microtubule organization, dynamics, and/or posttranslational modification status has been shown to have significant effects on the migration of multiple corneal cell types both *in vitro* and *in vivo* [18–22]. Furthermore, arthritis patients treated with a drug that broadly depolymerizes microtubules display significantly reduced corneal wound healing. These data highlight both the promise of targeting microtubules to regulate wound healing and the need for a better understanding of how microtubule dynamics affect cell migration [23, 24].

Our previous work identified Fidgetin-like 2 (FL2) as a microtubule regulator important for cell migration. Specifically, FL2 localizes to the cell edge where it selectively severs dynamic microtubules to inhibit directional cell migration. As a result, silencing FL2 with small interfering RNA (FL2-siRNA) promotes cell motility *in vitro* and wound healing in animal models, such as dermal excision and burn wounds in mice [12, 13].

MT severing enzymes, which are members of the <u>A</u>TPases <u>A</u>ssociated with diverse cellular <u>A</u>ctivities (AAA+) superfamily, cause breakages in MTs by forming hexameric rings around the C-terminal tails of tubulin and using energy from ATP hydrolysis to pull on the tails, thereby causing tubulin dimers to dissociate from the MT lattice [25, 26]. Through their severing activity, they regulate MT length, number, and branching, and fine-tune the dynamics of the MT cytoskeleton [27]. MT severing enzymes include katanin, spastin, and the fidgetin family (fidgetin, fidgetin-like 1 (FL1), and fidgetin-like 2 (FL2)). FL2 is highly similar to canonical Fidgetin within its catalytic AAA domain but diverges elsewhere within its polypeptide sequence and is present only in vertebrates. Our previous work has demonstrated that silencing FL2 with siRNA promotes cell motility *in vitro* and wound closure in animal models [12, 13]. Here, using a dose-response study, we report that FL2 siRNA delivered via nanoparticle technology (FL2-NPsi) can promote the repair of alkali burned corneas and reduce corneal tissue edema and scarring within two weeks of injury.

## Methods

The study protocol was approved by the Albert Einstein College of Medicine IACUC, and was conducted in adherence to the ARVO Statement for the Use of Animals in Ophthalmic and Vision Research.

Eighty 12-week old, male Sprague-Dawley rats were divided evenly into eight groups: positive control prednisolone (prednisolone acetate, Pred Forte, Allergan, Irvine, CA), empty nanoparticles, control-NPsi (1 μM, 10 μM, 20 μM) and FL2-NPsi (1 μM, 10 μM, 20 μM), with 10 rats per group. After the animals were anesthetized with isoflurane gas and an injection of ketamine and xylazine, topical 0.5% proparacaine was applied to the right cornea before surgery. The corneal epithelium was removed with a foam tip applicator, and a chemical injury was induced on the right cornea of each rat using 4mm discs of 1M NaOH-soaked filter paper, applied directly for 10 seconds. Next, the eye was washed with 3 consecutive 10mL washes of sterile PBS. The left eye was left uninjured and untreated to serve as a negative control. The injured eye was then treated for the subsequent 14 days according to their group assignment described above, along with an antibiotic eye drop (0.3% ofloxacin, Allergan, Irvine, CA), with drops administered at least 10 minutes apart from each other to prevent washout. Nanoparticles were given every other day, as siRNA has an intracellular half-life of 24-72 hours [28], while prednisone treatments were given twice daily and antibiotic eye drops administered daily. Prednisolone acetate is known to have a half-life of 30 minutes in aqueous humor and is usually dosed at two to four times [29], while ofloxacin has a half-life of several hours, with significant concentrations still present 6-24 hours after administration [30].

### Clinical analysis

Clinical assessment of wound healing was conducted in a blinded fashion every other day throughout the treatment period by recording digital images of corneas with a Leica EZ4 Stereo Dissecting Microscope (10447197, Heerbrugg, Switzerland) on Days 0, 2, 4, 6, 8, 10, 12, and 14. After documenting the appearance of the corneas under normal lighting, the surface was stained with a drop of fluorescein solution (Fluress, Akorn Incorporated, Decatur, IL, USA), enabling visualization of the corneal epithelial defect. The areas of epithelial defect were quantified using ImageJ software and subsequently expressed as a percentage of the total corneal area. The results were recorded and assessed using the student’s t-test and ANOVA (N=6 per group). Additionally, the degree of corneal opacity was scored, noting presence and degrees of hemorrhage, edema, and neovascularization (**Supplemental 1**).

### Histopathologic analysis

At the end of the treatment period, animals were euthanized and eyes enucleated for histopathologic evaluation (N=6 per group). Corneal tissue was embedded in paraffin, stained with hematoxylin & eosin (H&E), and cut to a thickness of 5-7 μm. Analyses of the tissue specimens were conducted in a blinded fashion. To carry out the immunohistochemical analyses of the corneal tissues, 4 animals per group were sacrificed at 4 different time points: days 1, 3, 7, and 14. This timeline was selected because inflammatory cells such as neutrophils and macrophages are most prominent between days 2 and 4. The following antibodies were selected and tested on half of each cornea: anti-CD31 mouse monoclonal IgG1 targeting vascular endothelial cells (BioRad, Kidlington, Oxford, England), anti-Neutrophil rabbit polyclonal IgG targeting neutrophils (LifeSpan BioSciences, Seattle, WA, USA), and OX42 mouse monoclonal IgG2 targeting macrophages (Santa Cruz Biotechnology, Dallas, TX, USA). All antibodies were unconjugated and stained with a secondary, dye-conjugated antibody. Specimens were embedded in paraffin and cut to a thickness of 8-10 μm. Analysis of these specimen likewise occurred in a blinded manner.

### Toxicity analysis

Evaluation for toxicity was performed to assess for apoptosis and nerve damage. To assess for apoptosis, 9 male Sprague-Dawley rats, one rat per group (uninjured/untreated, prednisone, empty NP, control-NPsi at 1 μM, 10 μM, or 20 μM, FL2-NPsi at 1 μM, 10 μM, 20 μM), were wounded and treated as described previously. At the end of treatment, animals were euthanized and eyes were collected for analysis. Corneas were trimmed around the sclerolimbal region and fixed in 4% paraformaldehyde overnight. The transferase dUTP Nick End Labeling (TUNEL) assay was performed on cryosections of rat eyes using the TUNEL apoptosis detection kit (DeadEnd Fluorometric TUNEL System; Promega, Madison, WI) according to the manufacturer’s instructions. Corneal toxicity from each treatment was evaluated by determining TUNEL-positive cell density (number per square millimeter) calculated as follows: number of TUNEL-positive cells/total number of cells. Three random fields of view were taken per corneal tissue sample.

To assess for nerve damage, the corneas were stained with neuronal specific rabbit polyclonal BIII tubulin antibody (BioLegend, San Diego, CA). Corneas were flat mounted on slides and imaged using an EVOS FL Auto Imaging System (ThermoFisher, Waltham, MA). Innervation of the cornea was defined as tortuous nerve endings organized in a clustered pattern originating from a single larger nerve and extending in three dimensions. Clusters were identified and counted manually from epifluorescence images of intact and hemisected corneal whole mounts [31].

### siRNA nanoparticle synthesis

Nanoparticles were prepared as follows [12]: 500 μl of Tetramethyl orthosilicate (TMOS) was hydrolyzed in the presence of 100 μl of 1 mM HCl by sonication on ice for about 15 min, until a single phase formed. The hydrolyzed TMOS (100 μl) was added to 900 μl of 10 μM of pooled siRNA against rat FL2 (siRNA from Sigma-Aldrich (St Louis, MO): SASI_Rn02 00314854; SASI_Rn02 00314854; SASI_Rn02_00389576) or the negative control siRNA (Sigma, Universal Negative control B) solution containing 10 mM phosphate, pH 7.4. A gel was formed within 10 minutes. The gel was frozen at −80°C for 15 minutes and lyophilized. The dried sample was ground into a fine powder with a mortar and pestle. The nanoparticles were then resuspended in sterile PBS at an siRNA concentration of 1, 10, and 20 μM, and stored at −80 C until immediately before use.

### qPCR

For cell culture experiments, RNA was extracted with Trizol (Fisher, Hampton, NH) by standard protocol. 200-300 ng of RNA was reverse transcribed using the SSVilo IV kit (Invitrogen, Carlsbad, CA). PowerSYBR Green Master Mix (Invitrogen, Carlsbad, CA) was used for qPCR, using the 7300 Real-Time PCR system (Applied Biosystems); rat *Fignl2:* GAGTTGCTGCAGTGTGAATG and CTCTGTGCTTCTGTCTCTGT; rat *β–actin*: CGTTGACATCCGTAAAGACC (sense), TCTCCTTCTGCATCCTGTCA (antisense). Results were quantified using the comparative 2^-ΔΔCt^ method. For B35 cells, experiment was performed three times.

### Statistical analysis

All data analysis in this study was carried out using GraphPad Prism 6 or Microsoft Excel (version 16.16.18). The samples and animals were allocated to experimental groups and processed randomly. All *in vitro* experiments represented multiple independent experiments conducted in triplicate. The *in vivo* experiments were performed with 2 to 6 rats for each condition. All data are represented as means ± SEM. Statistical analyses for the above characteristics were performed using the unpaired Student’s t-test (for comparing two distributions) and one-way ANOVA for more than two distribution. Differences were considered statistically significant at a p-value of <0.05.

## Results

Alkaline chemical injuries were induced onto the right cornea of rats, which were subsequently treated with prednisolone, empty nanoparticles, control-NPsi (1 μM, 10 μM, 20 μM) and FL2-NPsi (1 μM, 10 μM, 20 μM). FL2-NPsi-treated groups demonstrated significantly greater corneal re-epithelialization rates when compared to the prednisone and their respective control concentration groups (**Figure 1 & Table 1**). Specifically, 10 and 20 μM FL2-NPsi were determined to be the most efficacious concentrations in reducing time to corneal re-epithelialization throughout the healing process.

**Figure 1.**
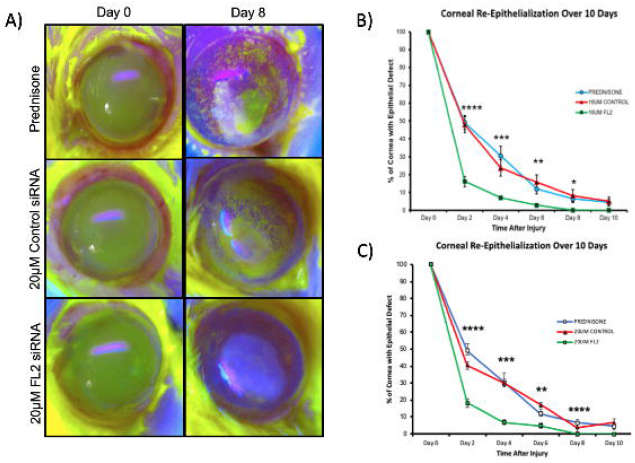
siRNA-Mediated Depletion of FL2 Enhances Corneal Re-Epithelialization. (A) Images of fluorescein-stained eyes eight days after injury. (B) Plots showing the kinetics of corneal re-epithelialization over 10 days with prednisone (blue line), 10 μM control-NPsi (red line), 10 μM FL2-NPsi (green line). (C) Plots showing the kinetics of corneal re-epithelialization over 10 days with prednisone (blue line), 20 μM control-NPsi (red line), 20 μM FL2-NPsi (green line). Data were pooled from multiple independent assays from corneas with alkaline injuries. The data for every time point were assessed using one-way ANOVA and the levels of significance shown (****=p<0.0001; ***=p<0.001; **=p<0.01; *=p<0.05; n.s., not significant).

**Table 1.** Percentages of Corneal Surface Area with Epithelial Defect Over 14 Days After Alkaline Injury.

| Percentage of Cornea Surface Area with Epithelial Defect |  |  |  |  |  |  |  |  |
| --- | --- | --- | --- | --- | --- | --- | --- | --- |
| Treatment Group | Day 0 | Day 2 | Day 4 | Day 6 | Day 8 | Day 10 | Day 12 | Day 14 |
| PREDNISONE | 100% | 49.2% | 30.4% | 11.9% | 6.5% | 4.4% | 0% | 0% |
| EMPTY NP | 100% | 39.1% | 21.8% | 17.0% | 7.5% | 4.6% | 0% | 0% |
| 1UM CONTROL NP-si | 100% | 36.5% | 22.5% | 12.4% | 9.7% | 3.9% | 0% | 0% |
| 10UM CONTROL NP-si | 100% | 48.0% | 23.7% | 15.7% | 8.2% | 5.1% | 0% | 0% |
| 20UM CONTROL NP-si | 100% | 40.5% | 29.9% | 17.1% | 13.7% | 6.4% | 0% | 0% |
| 1UM FL2 NP-si | 100% | 26.8% | 12.5% | 9.8% | 3.1% | 0% | 0% | 0% |
| 10UM FL2 NP-si | 100% | 16.2% | 7.0% | 2.7% | 0.1% | 0% | 0% | 0% |
| 20UM FL2 NP-si | 100% | 18.2% | 6.9% | 4.9% | 0% | 0% | 0% | 0% |
| 50UM FL2 NP-si | 100% | 31.7% | 11.2% | 9.0% | 7.3% | 2.4% | 0% | 0% |

The degree of corneal opacity was scored from a scale of 0-4 (**Supplemental 1**). The prednisone, empty NP, 1uM control-NPsi, 10uM control-NPsi, 20uM control-NPsi groups scored between 3 and 4 across all eyes. The 1 μM, 10 μM FL2-NPsi and 20 μM FL2-NPsi groups, however, scored between 1 and 2. When comparing control and FL2 treatment groups by concentration, the 20 μM FL2-NPsi-treated eyes demonstrated significantly decreased opacity with less peripheral neovascularization and central edema when compared to those treated with 20 μM of control-NPsi (**Figure 2**). However, the 10 μM and 1 μM FL2-NPsi groups were not significantly different from their control-NPsi counterparts.

**Figure 2.**
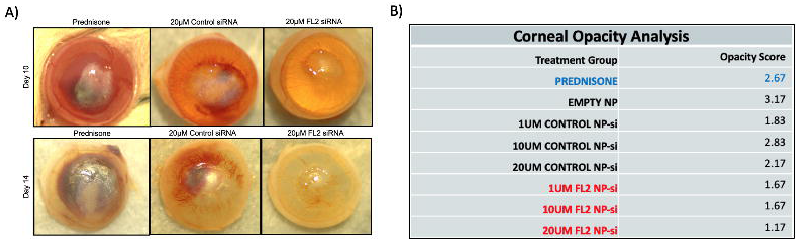
siRNA-Mediated Depletion of FL2 Enhances Corneal Transparency in Corneal Tissue. (A) Corneal appearance at Day 10 and Day 14. In the prednisone-treated group, corneal tissue is cloudy with poor central corneal healing after alkali injury. Peripheral neovascularization is extensive. In the 20 μM control-NPsi-treated group, corneal tissue is cloudy and exhibits extensive peripheral neovascularization. Some hemorrhage is seen behind the corneal tissue. In the 20 μM FL2-NPsi-treated group, the corneal tissue is more transparent and less edematous with less peripheral neovascularization. Central corneal tissue healing appears improved compared to prednisone and control-NPsi-treated groups. (B) Table of corneal opacity scores after 14 days.

After 14 days, all layers of corneal tissue were compared in a blinded fashion, including for epithelial healing (structure and organization), arrangement of stromal collagen lamellae, and presence of edema, inflammatory cells, and vascularization. Histopathologic analysis by a blinded ocular pathologist using a scoring system of 0-4+, with 0 being absent and 4+ being most severe, revealed 2-3+ stromal edema, inflammation, and neovascularization and 2+ retrocorneal membrane in control groups. In contrast, the FL2-NPsi-treated population exhibited fewer inflammatory cells and less neovascularization, with stromal edema, inflammation, and neovascularization scored as 1-2+ in the absence of retrocorneal membranes. The 10 μM FL2-NPsi group exhibited 2+ stromal edema, inflammation, and neovascularization but absence of anterior chamber inflammation, suggesting mild improvement compared to the aforementioned groups. Still, it was less efficacious than the 20 μM FL2-NPsi group, which yielded 1-2+ stromal edema, inflammation, and neovascularization with absence of anterior chamber inflammation (**Figure 3**).

**Figure 3.**
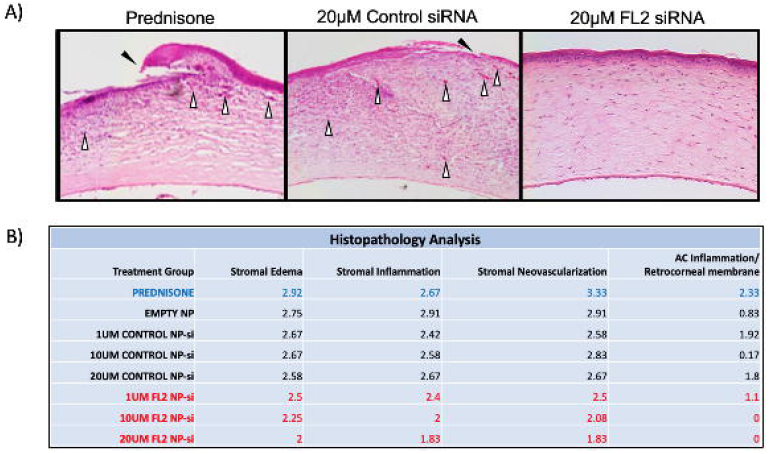
FL2-NPsi Treatment Stimulates Healing in the Rat Alkali Burn Injury Model. (A) Following 14 days of treatment, sections of corneal tissues were stained with H&E and examined. Representative images shown for rats treated with Prednisone, 20 μM control-NPsi, or 20 μM FL2-NPsi. The corneal epithelium layer is poorly healed, thinned, and irregular with superficial keratinization (black arrowheads) in prednisone and control-NPsi-treated groups. The corneal stroma is extremely edematous, and collagen lamella is disorganized and infiltrated with many inflammatory cells and extensive neovascularization (white arrowheads). In the FL2-NPsi-treated group, the corneal edema is less pronounced and the epithelium is thicker and more regular compared to prednisone or control-NPsi-treated groups. The corneal stroma shows less inflammatory cell infiltration and less neovascularization. Data is representative of ≥3 independent experiments. (B) Table of histopathologic scores after 14 days.

As there is less tissue in the wounded zone, confirming knockdown *in vivo* proved to be technically difficult. There is no reliable antibody against the rat homolog of FL2, so Western blotting to confirm KD was not an option. For these reasons and despite multiple attempts, confirming knockdown *in vivo* was not achieved. FL2-NPsi induced knockdown was however confirmed *in vitro* using B35 rat neuroblastoma cells (**Supplemental 2**).

Lastly, preliminary toxicology studies using quantitative microscopy suggest that alkaline chemical injuries are deleterious to corneal nerves, but FL2-NPsi is not. In fact, the 10 μM and 20 μM FL2-NPsi-treated groups had a greater number of nerve clusters as compared to control and untreated groups, demonstrating a similar profile to the uninjured/untreated negative control cornea (**Figure 4**), while empty nanoparticle and control-NPsi-treated groups demonstrated significantly reduced nerve densities. In addition, there was not a statistically significant difference in the number of apoptotic cells between prednisone, control, or FL2-NPsi-treated groups on the TUNEL assay (p=0.917; **Figure 5**), suggesting that FL2-NPsi is not toxic to corneal tissues.

**Figure 4.**
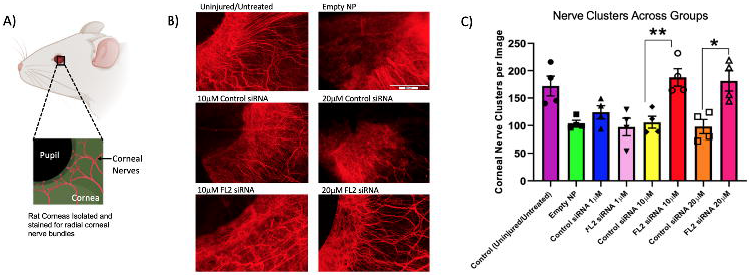
Corneal Nerve Assessment Using ß-III Tubulin Antibody. (A) Schematic of experiment for corneal nerve assessment. Corneal samples were derived from rats treated/untreated with siRNA NPs. Whole mount corneas were imaged after staining for corneal nerve bundles. (B) Corneas stained with ß-III tubulin antibody in corneal tissue reveal that empty NP and control-NPsi-treated eyes demonstrate similar nerve densities, while uninjured/untreated and FL2-NPsi-treated eyes exhibit comparable nerve densities. Red=ß-III Tubulin. (C) Quantification of nerve clusters across treatment groups reveal that the 10 μM and 20 μM FL2-NPsi-treated corneas demonstrate a greater number of nerve clusters as compared to control groups, and is comparable to that of uninjured/untreated corneas.

**Figure 5.**
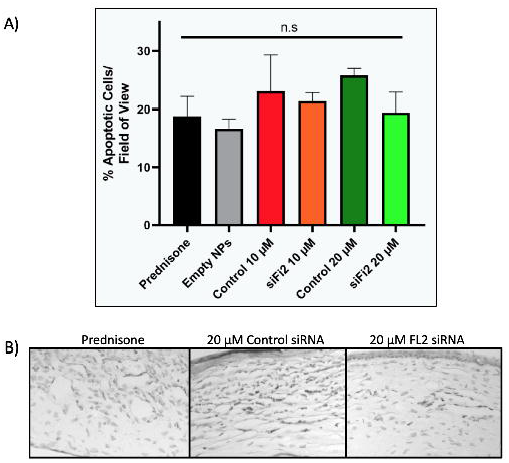
Apoptosis in Corneal Tissue Using TUNEL Assay. (A) Prednisone, control-NPsi, and FL2-NPsi-treated corneas exhibit no statistically significant difference in the number of apoptotic cells after alkaline chemical injury. n.s., not significant. (B) Apoptotic cells in prednisone, 20 μM control-NPsi, and 20 μM FL2-NPsi-treated corneas.

## Discussion

In recent years, there have been multiple investigations into harnessing the intracellular cytoskeleton, particularly microtubules, in tissue repair. Along with their structural role, microtubules are an important substrate for many of the molecular motor proteins responsible for intracellular transport. Regulation of the dynamic properties of microtubules are critically important (especially near the cell periphery) for the capacity of cells to close wounds. Our previous work screening biological agents for cell migration phenotypes identified a novel regulator of the microtubule cytoskeleton, termed Fidgetin-like 2 (FL2), as an important negative regulator of cell migration[12]. siRNA-mediated depletion of FL2 results in a significant increase in the velocity and directionality of cells *in vitro* [12]. Moreover, we found that *in vivo* depletion of FL2 strongly promoted wound closure and repair [12, 13]. FL2 depletion enhanced healing by inducing a targeted, transient disruption of cortical microtubules within the leading edge of cells [12]. Charafeddine *et al* (2015) translated these basic cell biology findings into an *in vivo* murine model of wound healing using a nanoparticle formulation to deliver siRNA to wounded skin. Here, we applied the same core nanoparticle technology to enhance corneal repair.

Exposure of the cornea to alkaline solutions, particularly ammonia, can result in near immediate damage to the corneal basement membrane, stromal keratocytes and nerve endings, endothelium, lens epithelium, and vascular endothelium of the conjunctiva, episclera, iris, and ciliary body. The penetration of ammonia can cause saponification of cell membranes leading to cell death, hydration of glycosaminoglycans leading to reduced stromal clarity, hydration of collagen fibrils leading to alterations of the trabecular meshwork, and the release of prostaglandins [32–34]. As a result, significant inflammation typically occurs and can dramatically slow the rate of epithelial cell migration into the wound zone [35–37].

Recovery towards an intact corneal epithelium is noted to be the most important determinant for a favorable outcome following chemical injury [4], since its presence signals a positive feedback loop that limits the production of collagenase in the stroma after re-epithelialization, thereby maintaining corneal transparency [38–40]. Studies have demonstrated that an intact epithelium may also be critical in preventing additional rounds of inflammation. When an epithelial defect remains after seven days, a second wave of inflammation begins and persists [41, 42]. Therefore, identifying a factor that can expedite the migration of cells to reform an intact corneal epithelium can dramatically reduce inflammatory complications and improve outcomes.

When considering corneal re-epithelization, the 10 μM and 20 μM FL2-NPsi groups demonstrated the greatest rates of epithelial healing. In contrast, the 1 μM FL2-NPsi-treated group performed worse than the 10 and 20 μM concentrations, most likely because 1 μM is too low a concentration to produce a significant effect on gene expression. Still, all FL2 NPsi-treated groups showed a greater rate of re-epithelialization compared to control-NPsi, empty NP, and prednisone-treated groups (**Figure 1 & Table 1**).

In addition to efficacy, the safety of FL2-NPsi is an important consideration for clinical translation. Interestingly, our preliminary toxicology evaluation using quantitative microscopy to assess for nerve damage found that FL2-treated corneas exhibited similar profiles to uninjured/untreated negative controls. This finding suggests that FL2 may not only be non-toxic to corneal nerves, but may also play a role in nerve restoration after alkaline chemical injury. In fact, a recent study by Baker *et al*., found that FL2 knockdown had a neuro-restorative effect on injured cavernous nerves in a rodent model [43]. The increase in the number of nerve clusters observed in the 10 μM and 20 μM of FL2-NPsi groups warrants further studies to assess for the possible neuro-restorative potential of this treatment (**Figure 4**).

The current use of siRNA to treat ocular maladies presents an area ripe for future development. Because the eye is a confined compartment and easily accessible for topical treatment, siRNA may be particularly promising [44]. Recent approaches using siRNAs to down-regulate the expression of genes associated with proliferation, fibrosis, or inflammation (e.g. CAP37, Caveolin-1, TGFB1, TGFBR2, CTGF, VEGFR1) have shown a range of promising preclinical data. However, these targeted proteins are involved in known cancer pathways, limiting clinical translation and underscoring the need to identify a new gene target [45–48].

In summary, rapid recruitment of epithelial cells responsible for closing wounds and stabilizing the corneal surface may be key in wound healing. By enhancing cell motility, wound healing can occur more rapidly and with high fidelity to the original tissue. This can have several profound effects on recovery, including reduced scarring, pain, risk of infection, and improved vision and restoration of corneal architecture.

## Acknowledgements

The authors would like to acknowledge Peng Guo, PhD for his help with analytical imaging.

**Supplemental 1. Clinical Assessment of Corneal Opacity – Scoring Sheet.**

**Supplemental 2. Nanoparticle-Mediated Knockdown of FL2 in an In Vitro qPCR Assay.** A significant knockdown (49.6% of control) of FL2 was detected, illustrating the ability of the NPs to successfully deliver siRNA to cells.

