## Supplemental 1 for "“A Novel Therapeutic Approach to Corneal Alkaline Burn Model by Targeting Fidgetin-like 2, a Microtubule Regulator”"

Clinical Assessment of Corneal Opacity – Scoring Sheet

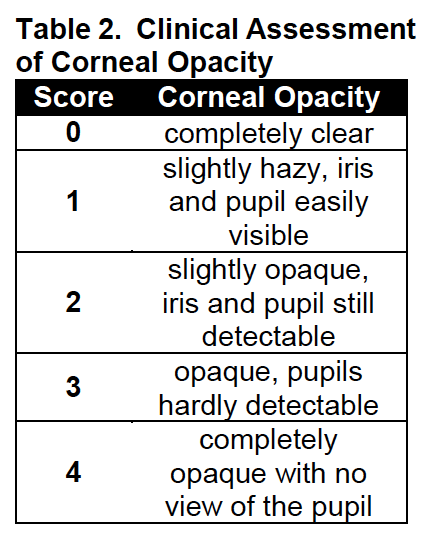

Picture ID: _______

1) **Corneal Opacity: _____**

Example:

Grade 0	Grade 1	Grade 2	Grade 3	Grade 4

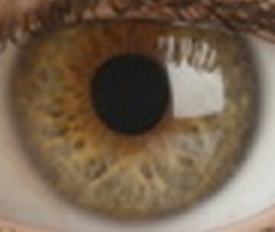

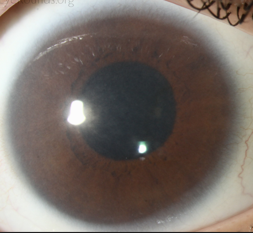

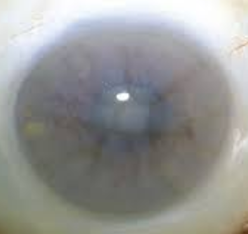

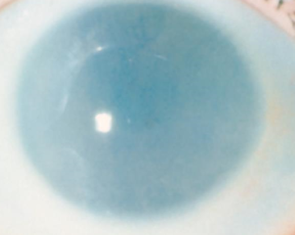

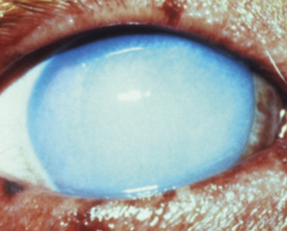

2) **Please note clinical signs of the following, if present:**
a) **Hemorrhage:**

**________________________________________________________________________________________________________________________________________________**

b) **Edema:**

**________________________________________________________________________________________________________________________________________________**

c) **Peripheral Corneal Neovascularization:**

**________________________________________________________________________________________________________________________________________________**
