## Supplementary figures and images for "“A Novel Therapeutic Approach to Corneal Alkaline Burn Model by Targeting Fidgetin-like 2, a Microtubule Regulator”"

### Supplemental 2

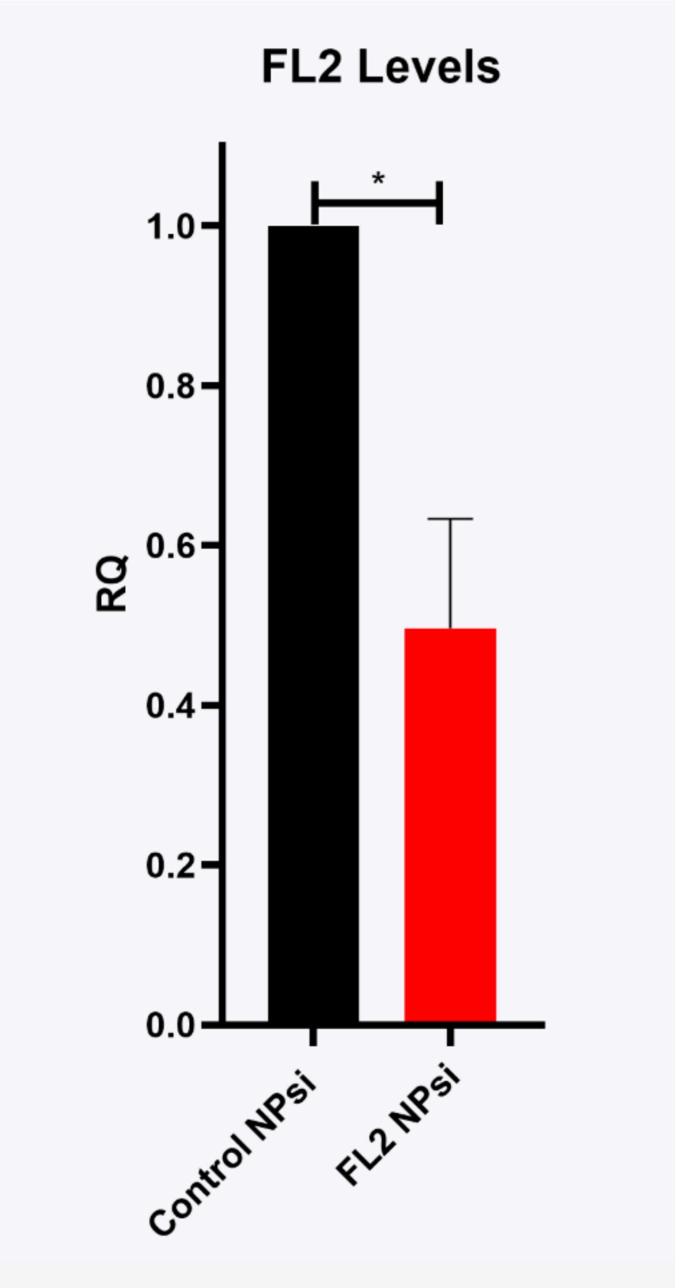
